# Extended field-of-view ultrathin microendoscopes with built-in aberration correction for high-resolution imaging with minimal invasiveness

**DOI:** 10.1101/504472

**Authors:** Andrea Antonini, Serena Bovetti, Claudio Moretti, Francesca Succol, Vijayakumar P. Rajamanickam, Andrea Bertoncini, Carlo Liberale, Tommaso Fellin

## Abstract

We present a novel approach to correct optical aberrations in ultrathin gradient-index rod lens-based endoscopes using microfabricated aspherical lenses. Corrected microendoscopes have up to 9 folds larger field-of-view compared to uncorrected probes. Using extended field-of-view (*eFOV*) microendoscopes, we report two-photon imaging of GCaMP6 signals in the mouse hippocampus *in vivo* with unprecedented combination of high spatiotemporal resolution and minimal invasiveness.

## Introduction

Two-photon fluorescence imaging allows high resolution anatomical and functional visualization of neuronal circuits several hundred of micrometers deep into the intact mammalian brain (1). Light scattering within the brain, however, strongly affects the propagation of excitation and emission photons, making effective imaging increasingly difficult with tissue depth (2, 3). Various strategies have been developed to improve imaging depth in multi-photon fluorescence microscopy (4–11), allowing the visualization of regions 1–1.6 mm below the brain surface. However, deeper imaging requires the use of implantable microendoscopic probes, which allow optical investigation of neural circuits in brain regions that would otherwise remain inaccessible (12–16). Ideally, microendoscopic devices should have small radial dimensions and, at the same time, maintain sub-cellular resolution across a large field-of-view (FOV). This would allow high-resolution population imaging, while minimizing tissue damage. Current microendoscopes for deep imaging are frequently based on gradient-index (GRIN) rod lenses which typically have diameter between 0.35–1.5 mm and are characterized by intrinsic optical aberrations (14). These aberrations are detrimental in two-photon imaging because they decrease the spatial resolution and lower the excitation efficiency, leading to degraded image quality and restricted FOV (17, 18). This is especially relevant when ultrathin microendoscopes (diameter ≤ 500 μm) are used, because the size of the FOV decreases with the diameter of the optical probe. Optical aberrations in GRIN microendoscopes can be corrected with adaptive optics which, however, requires significant modification of the optical path (17, 19, 20) and may limit the temporal resolution of functional imaging over large FOVs (17). Alternatively, the combination of GRIN lenses of specific design with plano-convex lenses within the same microendoscopic probe has been used to increase the Numerical Aperture (NA) and to correct for aberrations on the optical axis (14). However, technical limitations in manufacturing high-precision optics with small lateral dimensions have so far prevented improvements in the performances of GRIN microendoscopes with lateral diameter < 1 mm using corrective optical microelements (21). Here we report the development and application of a new approach to correct aberrations and extend the FOV in ultrathin GRIN-based endoscopes using microfabricated aspheric lenses. This method has no limitations on the size of the corrective lenses for current GRIN-based microendoscopes and requires no modification of the microscope optical path. Corrective lenses were optically designed based on ray trace simulations and microfabricated using two-photon polymerization (TPP) (22). We developed four types of extended field-of-view (*eFOV*) ultrathin microendoscopes, differing in length and diameter, which enable optimized optical performances at various depths within biological tissue. Finally, we validated *eFOV*-microendoscopes performing functional imaging on hundreds of hippocampal cells expressing the genetically-encoded calcium indicator GCaMP6 (23) in the intact mouse brain *in vivo* with an unprecedented combination of high resolution, FOV extension and minimal invasiveness.

## Materials and Methods

### Optical design and simulation

Simulations were run with OpticStudio15 (Zemax, Kirkland, WA) to define the profile of the aspheric corrective lens to be integrated in the *eFOV*-microendoscopes, with the aim to achieve: *i*) a full-width half maximum (FWHM) lateral resolution < 1 μm at the center of the FOV; *ii*) a FWHM axial resolution below < 10 μm; *iii*) a working distance between 150 μm and 220 μm into living brain tissue. The wavelength used for simulations was λ = 920 nm since the devices were developed for two-photon functional microscopy applications using GCaMP6s (23).

The surface profile of corrective aspheric lenses has been described as (24):

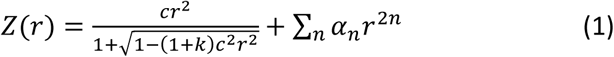

Since GRIN lenses have intrinsic spherical aberration, the optimization for the shape of the corrective lenses started with the profile of a Schmidt corrector plate (25) as initial guess; the parameters c, k, α_n_ (with n = 1–8) in equation (1) were then automatically varied in order to maximize the Strehl ratio (26) over the largest possible area of the FOV (Supplementary table 1). A fine manual tuning of the parameters was performed for final optimization.

### Corrective lens manufacturing and endoscope assembly

The optimized aspheric lens structure obtained with simulations was exported into a 3D mesh processing software and converted into a point cloud dataset fitting the lens surface (with ~ 300 nm distance among first neighborhood points). Two-photon polymerization with a custom set-up (22) including a dry semi-apochromatic microscope objective (LUCPlanFLN 60x, NA 0.7, Olympus Corp., Tokyo, JP) and a near infrared pulsed laser beam (duration, 100 fs; repetition rate, 80 MHz; wavelength, 780 nm; FemtoFiber pro NIR, Toptica Photonics, Graefelfing, DE) was used for the fabrication of the corrective lenses. A drop of resin (4,4ʹ-Bis(diethylamino)benzophenone photoinitiator mixed with a diacrylate monomer), sealed between two coverslips, was moved by a piezo-controlled stage (model P-563.3CD, PI GmbH, Karlsruhe, DE) with respect to the fixed laser beam focus, according to the 3D coordinates of the previously determined point cloud, with precision of 20 nm. Output laser power was ~ 15 mW at the sample. Once the surface was polymerized, the lens was dipped for ~ 2 minutes in methanol followed by ~ 1 minute immersion in isopropyl alcohol, and finally exposed to UV light (λ = 365 nm; 3 Joule / cm^2^) to fully polymerize the bulk of the structure. In the second stage of the project, a commercial TPP fabrication system (Photonic Professional GT, Nanoscribe GmbH, DE) was also used for corrective lens fabrication.

For fast generation of multiple lens replicas, a molding (27) technique was used. To this end, polydimethylsiloxane (PDMS, Sylgard 164, 10:1 A:B, Dow Corning, Auburn, MI) was casted onto the lens and hardened by heat cure in a circulating oven at 80°C for approximately 30 minutes. The resulting bulked structure of solid PDMS was then used as negative mold. A drop of a UV-curable optically-clear adhesive with low fluorescent emissivity (NOA63, Norland Products Inc., Cranbury, NJ) was deposited on the negative mold, pressured against a coverslip (5 mm diameter) of appropriate thickness (100–200 μm thick depending on the *eFOV*-microendoscope type, Fig. 1) and hardened by UV exposure. The coverglass with the lens attached was detached from the mold and glued onto a metal ring. One end of the appropriate GRIN rod (NEM-050-25-10-860-S; NEM-050-43-00-810-S-1.0p; GT-IFRL-035-cus-50-NC; NEM-035-16air-10-810-S-1.0p, all purchased from Grintech GmbH, Jena, DE) was attached perpendicularly to the coverslip surface using NOA63. Alignment of the corrective lens and the GRIN rod was performed under visual guidance using an opto-mechanical stage (Supplementary Fig. 1a). An additional and removable coverglass (No. 1.5) was glued on the top of every support ring (Supplementary Fig. 1d) to keep the polymeric corrective lens clean and to protect it from mechanical damage. The GRIN rods were finally coated with a thin (< 30 nm) layer of polytetrafluoroethene using reactive ion etching.

**Figure 1.**
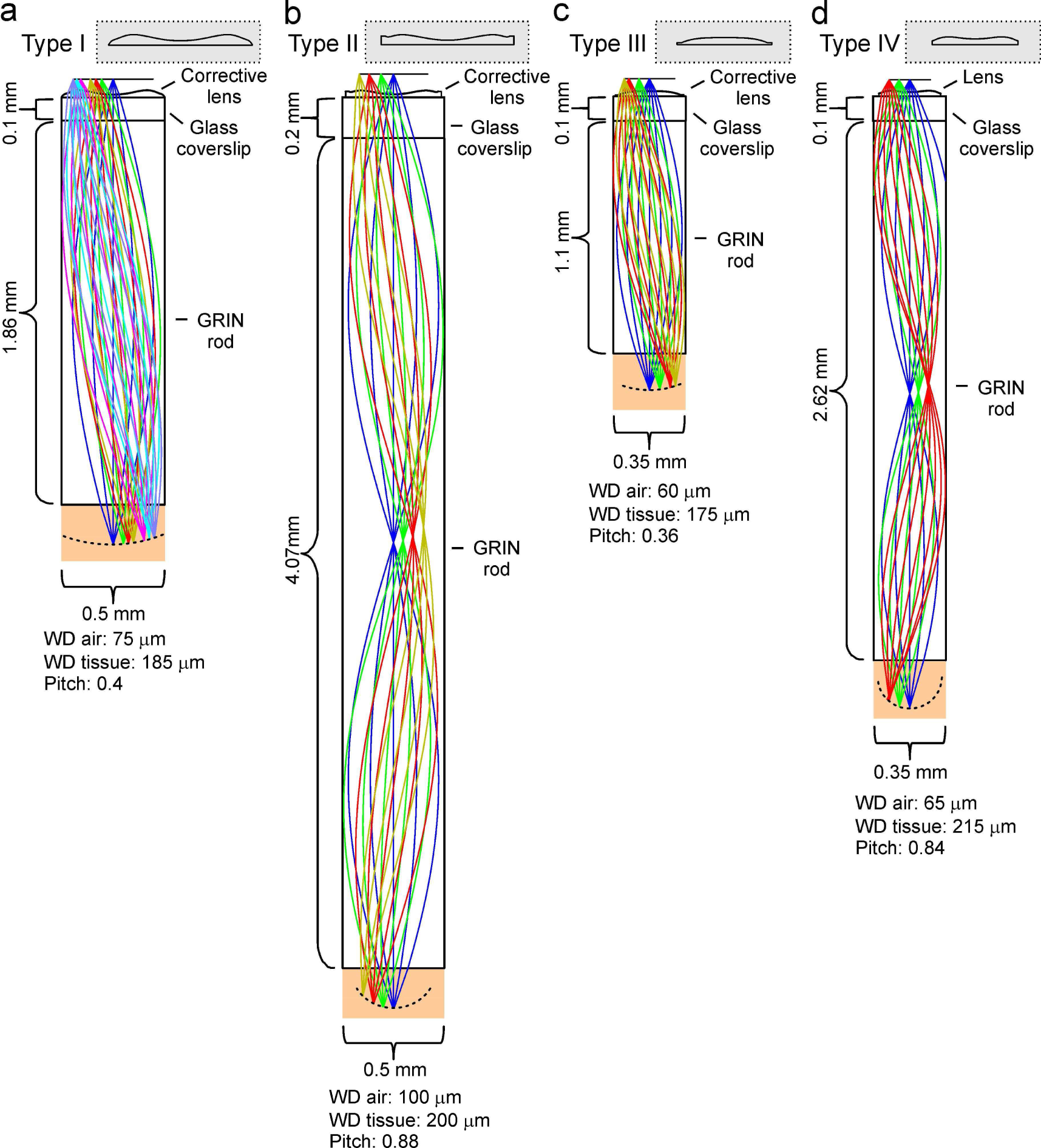
Optical design of *eFOV*-microendoscopes. **a-d)** Images showing ray trace simulations for the four different *eFOV*-microendoscopes (type I-IV) developed in this study. The insets show the profiles of corrective polymeric lenses used in the different *eFOV*-microendoscopes.

### Optical characterization

Optical characterization of *eFOV*-microendoscopes was carried out with a two-photon laser-scanning microscope equipped with a wavelength-tunable, ultrashort-pulsed, mode-locked Ti:Sapphire laser source (Ultra II Chameleon, pulse duration, 160 fs; repetition rate, 80 MHz; wavelength, 920 nm; Coherent Inc., Santa Clara, CA), a commercial Prairie Ultima IV scanhead (Bruker Corporation, Milan, IT, former Prairie Technologies) and an upright epi-fluorescence microscope (BX61 Olympus Corp., Tokyo, JP). For all measurements, the wavelength was set at 920 nm. The optomechanical assembly used for the microendoscope characterization is shown in Supplementary Fig. 1c. Two infinity-corrected objectives were used: RMS20X-PF-20X, and LUCPLFLN 60X, 0.7NA (Olympus Corp., Tokyo, JP). Measurement of maximal resolution was performed with the LUCPLFLN 60X objective to overfill the *eFOV*-microendoscope numerical aperture and reach diffraction limited performances in accordance with optical simulations, whereas other characterizations were done with the RMS20X-PF-20X objective. The maximal resolution of each *eFOV*-microendoscope was evaluated using subresolution spherical fluorescent beads (diameter: 200 nm, Polyscience, Warrington, PA), following a previous spatial calibration using a custom fluorescent ruler.

To evaluate the radial profile of the fluorescence intensity across the FOV and the extent of the aberration-corrected FOV, we used thin (thickness: 300 nm) fluorescent slices (28) and acquired z-series of images (512 pixels × 512 pixels) with 1 μm axial step. Image analysis was carried out following previous reports (28) using the ImageJ/Fiji software (29) and custom Python code. In x,z projections, we first fitted the fluorescence intensity profile with a circular section. We then averaged the fitting circular section across x,z projections obtained from different z-stacks for the same GRIN rod. We finally measured the mean curvature of the FOV of that specific GRIN rod from the average circular section. The axial projection of each z-stack was remapped onto the fitting circular section to optimally estimate radial distances.

For fluorescence intensity measurements, in each z-stack we measured the fluorescence intensity along *N* randomly chosen radial directions (*N* = 400). Fluorescence intensity along a given direction was initially smoothed with a ~7 μm flat moving window, averaged across *N* and normalized to the maximal intensity value. In these experiments, the diameter of the FOV was measured based on a threshold set at 80%. Intensity variations < 5% below the threshold were not considered if restricted to < 100 μm.

For measurements of the axial sectioning capability, at every x,y position the fluorescence intensity distribution along the direction perpendicular to endoscope FOV was fitted with a Gaussian curve. The FWHM was used as measurement of the z-resolution (FWHMz). For this analysis, the ImageJ/Fiji plug-in MetroloJ (30) and a Python code was used. We operationally defined the usable FOV as the area in which the FWHM_z_ remained < 1.5 × FWHM_z_, _min_, where FWHM_z_, _min_ is the FWHM_z_ at the center of the FOV.

### Animal surgery, viral injection and microendoscope implant

Experimental procedures involving animals have been approved by the Istituto Italiano di Tecnologia Animal Health Regulatory Committee, by the National Council on Animal Care of the Italian Ministry of Health (authorization # 1134/2015-PR) and carried out according to the guidelines of the European Communities Council Directive. Animals were housed under a 12-hour light:dark cycle in individually ventilated cages. Experiments were performed on adult (8–10 week old) C57BL/6J (Charles River, Calco, IT), and Scnn1a-Cre (B6;C3-Tg(Scnn1a-cre)3Aibs/J, Jackson Laboratory, Bar Harbor, USA) mice. Adeno-associated viruses (AAVs) AAV1.Syn.GCaMP6s.WPRE.SV40, AAV1.Syn.flex.GCaMP6s.WPRE.SV40, AAV1.CAG.Flex.eGFP.WPRE.bGH, AAV1.CaMKII0.4.Cre.SV40 were purchased from the University of Pennsylvania Viral Vector Core. Viral injections and *eFOV*-microendoscopes insertion were performed during a single surgical procedure. Animals were anesthetized with 2% isoflurane, placed into a stereotaxic apparatus (Stoelting Co, Wood Dale, IL) and maintained on a warm platform at 37°C. A small hole was drilled through the skull and 0.5 - 1 μl (30 nl/min, UltraMicroPump UMP3, WPI, Sarasota, FL) of AAVs containing solution was injected at stereotaxic coordinates: 1.4 mm posterior to bregma (P), 1 mm lateral to the sagittal sinus (L) and 1 mm deep (D) to target the hippocampal CA1 region; 1 mm anterior to bregma (A), 2 mm L and 2 mm D to target the dorsal striatum; 1.7 mm P, 1.6 mm L and 3 mm D to target the ventral posteromedial thalamic nucleus (VPM). Co-injection of AAV1.Syn.flex.GCaMP6s.WPRE.SV40 and AAV1.CaMKII0.4.Cre.SV40 (1:1) was performed to express GCaMP6s in hippocampus CA1 pyramidal cells. Injection of AAV1.Syn.GCaMP6s.WPRE.SV40 was performed to express GCaMP6s in the dorsal striatum. Injection of AAV1.Syn.flex.GCaMP6s.WPRE.SV40 in the Scnn1a-Cre mouse line was performed to express GCaMP6s in the VPM. Layer IV expression of GFP was achieved by injecting AAV1.CAG.Flex.eGFP.WPRE.bGH in the somatosensory cortical area of Scnn1a-Cre mice. Following virus injection a craniotomy (~ 600 × 600 μm^2^ or ~ 400 × 400 μm^2^ depending on the endoscope size) was performed over the neocortex at stereotaxic coordinates: 1.8 mm P and 1.5 mm L to image the hippocampus; 0.7 mm A and 1.8 mm L to reach the dorsal striatum; 2.3 mm P and 2 mm L to reach the VPM. A thin column of tissue was suctioned with a glass cannula (ID, 300 μm and OD, 500 μm; Vitrotubs, Vitrocom Inc., Mounting Lakes, NJ) glued onto the lateral side of transparent supports. The *eFOV*-microendoscope was slowly inserted in the cannula track, down to the depth of interest and secured by acrylic adhesive and dental cement to the skull. If necessary, metal spacers (thickness: ~ 100 μm) were glued on the flat coverslip surface to obtain the desired protrusion distance of the GRIN rod. After surgery, animals were positioned under a heat lamp and monitored until recovery. Three to five weeks after injection, mice were anesthetized with urethane (2 g/kg) and placed into a stereotaxic apparatus to proceed with imaging experiments. Body temperature was kept at 37 °C and depth of anesthesia was assured by monitoring respiration rate, eyelid reflex, vibrissae movements, and reactions to pinching the tail and toe. In some experiments, oxygen saturation was controlled by a pulseoxymeter (MouseOx, Starr Life Sciences Corp., Oakmont, PA).

### Functional imaging with eFOV microendoscopes in vivo

For scanning imaging of GCaMP6-expressing neurons, the same microscope set-up used for the optical characterization of *eFOV*-microendoscopes was used. GCaMP6 fluorescence was excited at 920 nm (laser power: 28–64 mW). For high-speed scanless imaging experiments, a liquid crystal spatial light modulator (SLM, X10468–07, Hamamatsu, Milan, IT) was placed in a plane optically conjugated with the galvanometric mirror aperture and detection of fluorescence signals was performed with a camera (SciMeasure NeuroCCD-SMQ, Redshirt Imaging, Decatur, GA), as previously described (31–33). Laser power *per* point was ~ 18 mW / point measured as the total power value under the endoscope divided by the number of projected points. Experiments were performed using an Olympus RMS20X-PF - 20X 0.5 NA Plan Fluorite Objective.

Temporal series recorded in the scanning configuration were imported into the open source ImageJ/Fiji (29) software and movement correction was performed using the plugin Image Stabilizer. Calcium traces were analyzed using a custom code based on the open-source CellSort MATLAB toolbox (34). Briefly, the motion-corrected image stack was normalized and analyzed by principal components analysis (PCA) to find and discard dimensions that mainly represented noise. Principal components displaying a variance greater than noise were then analyzed with an iterative independent component analysis (ICA) algorithm to identify active cells. Manual validation of extracted traces was performed. Signals *S_i_*(*t*) were standardized as 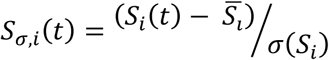
where 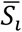 and σ(*S_i_*) are respectively the mean of the signal and its standard deviation (s.d.). For scanless imaging, t-series were imported into the open source ImageJ/Fiji software and the regions of interest (ROIs) were manually identified. The fluorescence signals corresponding to the ROIs were computed as *S_σ,i_*(*t*) as described above. In some cases, traces were filtered with an exponential weighted moving average (τ = 100 ms).

### Immunohistochemistry

Deeply anesthetized animals were transcardially perfused with 0.01 M PBS (pH 7.4) followed by 4 % paraformaldehyde. Brains were post-fixed for 6 h, cryoprotected with 30 % sucrose solution in 0.1 M PBS and serially cut in coronal sections (thickness: 40 - 50 μm). Sections were then counterstained with Hoechst (1:300, Sigma Aldrich, Milan, IT), mounted and coverslipped with a DABCO [1,4-diazobicyclo-(2,2,2)octane]-based antifade mounting medium. Fluorescence images were acquired with a Leica SP5 inverted confocal microscope (Leica Microsystems, Milan, IT).

### Statistics

Values are expressed as mean ± standard deviation, unless otherwise stated. A Kolmogorov-Smirnov normality test was run on each experimental sample. When comparing two populations of data, Student’s *t*-test was used to calculate statistical significance in case of Gaussian distribution.

## Results and Discussion

Four types (type I-IV) of *eFOV*-microendoscopes of various length and lateral dimensions were developed, all composed of a GRIN rod, a glass coverslip and a microfabricated corrective aspheric lens (Fig. 1). One end of the GRIN rod was directly in contact with the glass coverslip and the GRIN rod was different in each of the four types of *eFOV*-microendoscopes (lateral diameter, 0.35–0.5 mm; length, 1.1–4.1 mm; all 0.5 NA, table 1). The glass coverslip was 100 μm thick for type I, III, IV *eFOV*-microendoscopes and 200 μm thick for type II *eFOV*-microendoscopes. This design did not require additional cannulas or clamping devices (35, 36) that would increase the lateral size of the microendoscope assembly or reduce it usable length, respectively.

**Table 1.** Physical and optical characteristics of *eFOV*-microendoscopic probes. For statistical comparison of corrected *vs* uncorrected microendoscopes Student’s *t*-test was used.

| Type | Diam.<br>(mm) | Length<br>(mm) | FWHM<sub>x,y</sub><br>($\mu\text{m}$)<br>N = 10 | FWHM<sub>z</sub><br>($\mu\text{m}$)<br>N = 10 | FOV diam.<br>($\mu\text{m}$)<br>N = 7 |
| --- | --- | --- | --- | --- | --- |
| I | 0.5 | 1.86 | uncor.: $0.89 \pm 0.04$<br>cor.: $0.82 \pm 0.01$<br>p = 1.2E-4 | uncor.: $8.5 \pm 0.4$<br>cor.: $6.7 \pm 0.2$<br>p = 2.6E-10 | uncor.: $169 \pm 13$<br>cor.: $398 \pm 18$<br>p = 5.0E-12 |
| II | 0.5 | 4.07 | uncor.: $0.88 \pm 0.04$<br>cor.: $0.88 \pm 0.03$<br>p = 8.5E-1 | uncor.: $9.7 \pm 0.5$<br>cor.: $7.3 \pm 0.2$<br>p = 1.5E-9 | uncor.: $120 \pm 6$<br>cor.: $365 \pm 14$<br>p = 1.4E-14 |
| III | 0.35 | 1.1 | uncor.: $0.81 \pm 0.09$<br>cor.: $0.76 \pm 0.02$<br>p = 9.5E-2 | uncor.: $8.1 \pm 0.3$<br>cor.: $6.3 \pm 0.3$<br>p = 7.8E-11 | uncor.: $168 \pm 4$<br>cor.: $187 \pm 4$<br>p = 1.8E-5 |
| IV | 0.35 | 2.62 | uncor.: $0.82 \pm 0.05$<br>cor.: $0.77 \pm 0.08$<br>p = 9.7E-2 | uncor.: $8.0 \pm 0.4$<br>cor.: $6.4 \pm 0.5$<br>p = 4.8E-7 | uncor.: $133 \pm 2$<br>cor.: $233 \pm 10$<br>p = 1.1E-11 |

Corrective lenses were fabricated by TPP (22) and plastic molding replication (27) directly onto the glass coverslip. For each type of GRIN rod used in the *eFOV*-microendoscope, ray trace simulations determined the lens profile (Fig. 1) that corrected optical aberrations and maximized the FOV (Fig. 2). In the representative case of type I *eFOV*-microendoscope, the coefficients used in equation (1) were: c: −2.579E-1, k:-1.74, α_1_: 8.575E-1, α_2_:-5.297E1, α_3_: 5.952E3, α_4_: −2.765E5, α_5_: 7.258E6, α_6_: - 8.914E7, α_7_: 2.469E8, α_8_: 2.193E9. For type I *eFOV*-microendoscopes the corrective lens had a diameter of 0.5 mm and height < 40 μm. For this type of *eFOV*-microendoscopes, the simulated diffraction point-spread-functions (PSFs) at incremental radial distances (from 0 to 200 μm) from the optical axis showed that the Strehl ratio of the system was > 80% (diffraction-limited condition according to the Maréchal criterion (37)) at a distance up to ~ 165 μm from the optical axis with the corrective lens, while up to ~ 70 μm for the same optical system without the corrective lens (Fig. 2a), leading to an increase in the area of the diffraction-limited FOV of ~ 5 times. The coefficients used in equation (1) for all other types of *eFOV*-microendoscope are reported in Supplementary table 1. Fig. 2b-d reports the Strehl ratio for corrected and uncorrected type II-IV *eFOV*-microendoscopes. The enlargement of the area of the FOV was ~ 2–9 times for these other types of *eFOV*-microendoscopes, compared to microendoscopes without the corrective lens.

**Figure 2.**
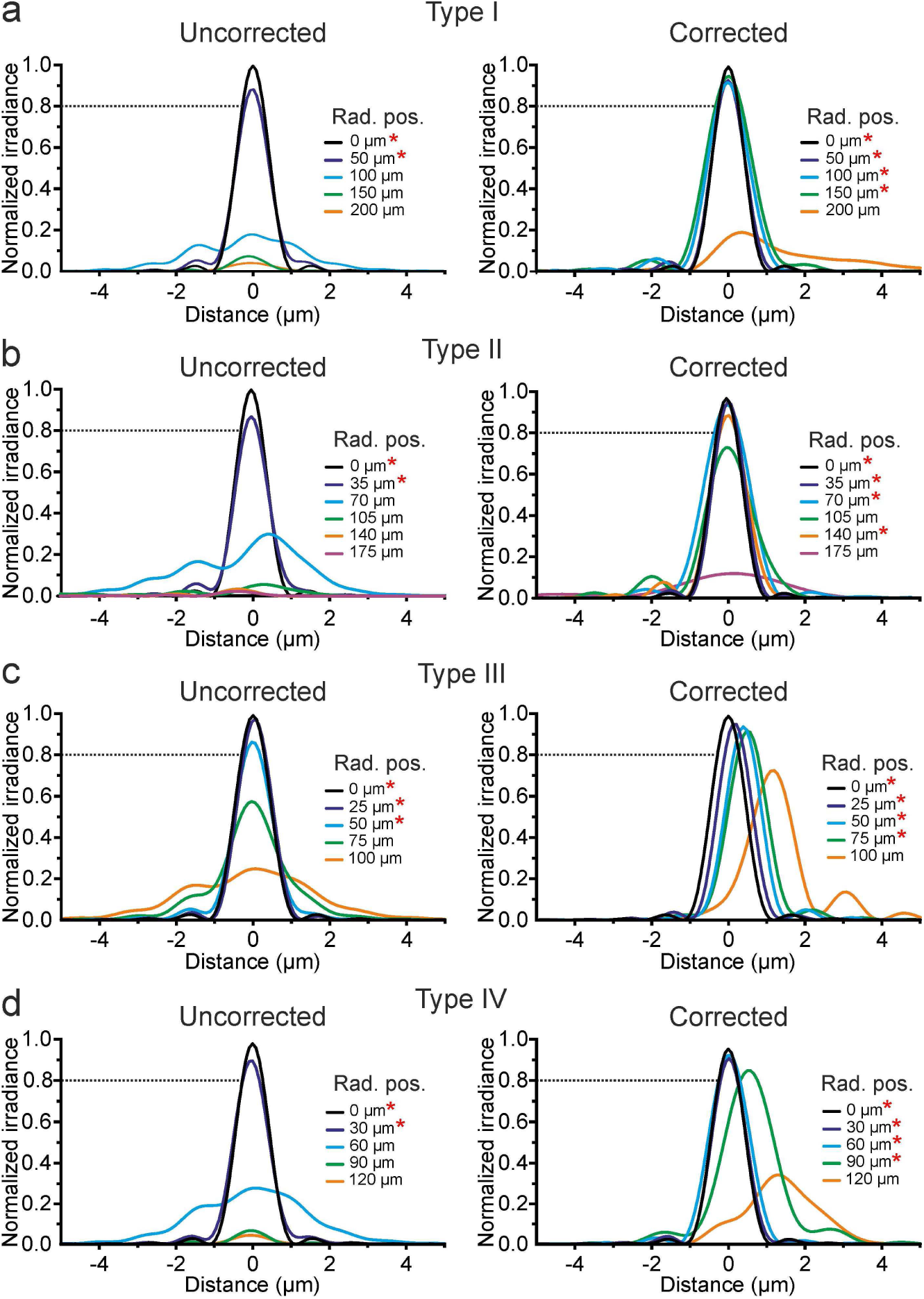
Corrective lenses improve the simulated optical performances of ultrathin microendoscopes. **a)** Simulated diffraction PSFs of type I microendoscopes without the corrective lens (uncorrected, left) and with the corrective lens (corrected, right). PSFs are shown color coded according to their radial positions (Rad. Pos.) from the optical axis. The black dotted line represents the diffraction-limited condition that was set at 80 % (Maréchal criterion). The red asterisks indicate the radial positions at which the maximal normalized irradiance of the corresponding PSF was > 80 %. **b-d)** Same as in a) for type II (b), type III (c), and type IV (d) microendoscopes.

To experimentally validate the optical performances of the *eFOV*-microendoscopes, we first coupled them with a standard two-photon laser scanning system using a customized mount (Fig. 3a-b, Supplementary Fig. 1). We measured the spatial resolution by imaging subresolution fluorescence beads (diameter: 200 nm) at 920 nm. We found that *eFOV*-microendoscopes had significantly improved axial resolution compared to uncorrected probes. For type I *eFOV*-microendoscopes, for example, the minimal value of FWHM_z_ was 6.7 ± 0.2 μm for corrected endoscopes and 8.5 ± 0.4 μm for uncorrected probes (Fig. 3d-e, table 1; Student’s *t*-test, p = 2.6E-10; N = 10). The radial resolution was also slightly increased for type I eFOV-microendoscopes (Table 1). Experimental measures of radial and axial resolution for type II-IV *eFOV-*microendoscopes are reported in Table 1. Importantly the axial resolution was significantly increased in all corrected *eFOV*-microendoscopes compared to uncorrected probes.

**Figure 3.**
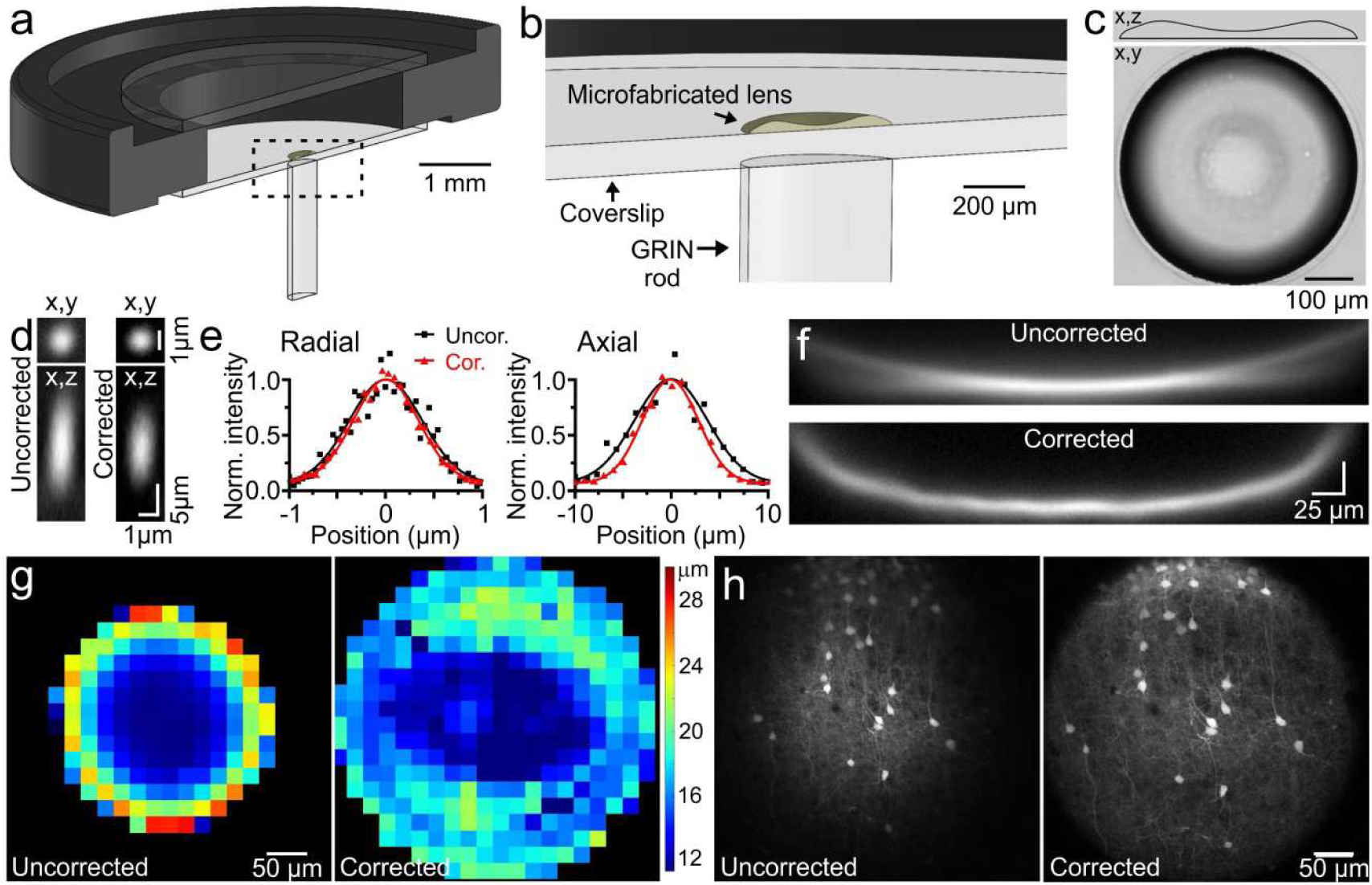
Aberration correction using microfabricated lenses extends the FOV in ultrathin microendoscopes. **a-b)** Schematic of the *eFOV*-microendoscope mount for head implant. The GRIN rod is attached to one side of the glass coverslip, the microfabricated polymeric lens to the other side of the coverslip. The coverslip is glued on a circular metal ring that facilitates fixation of the animal’s skull. The detail of the coupling between optical elements is shown at an expanded scale in b. **c)** Top: side profile of the corrective lens for type I *eFOV*-microendoscopes. Bottom: bright field image (top view) of the same corrective lens. **d)** Representative images in the x,y plane (top) and in the x,z plane (bottom) of a subresolution fluorescent bead (diameter: 0.2 μm) imaged with a type I *eFOV*-microendoscope without (uncorrected, left) and with (corrected, right) the microfabricated corrective lens. λ_exc_ = 920 nm. **e)** Normalized intensity profiles along the radial (left) and axial (right) directions of the images shown in d) for uncorrected (black) and corrected (red) type I microendoscopes. Data are normalized to the maximum of the Gaussian fit (solid line). **f)** x,z projections of a z-stack of two-photon laser scanning images of a subresolution fluorescent layer (thickness: 300 nm) obtained using a type I *eFOV*-microendoscope without (uncorrected, top) and with (corrected, bottom) the microfabricated corrective lens. **g)** Average axial resolution as a function of the position in the FOV for uncorrected (left) and corrected (right) type I microendoscopes. The pseudocolor scale indicates axial resolution values in microns. N = 7. **h)** Representative images of fixed cortical tissue expressing GFP in neuronal cells, acquired with the same microendoscope used in d without (uncorrected, left panel) and with (corrected, right panel) the microfabricated corrective lens.

We then evaluated the profile of fluorescence intensity across the FOV for both uncorrected and corrected probes using a subresolution thin fluorescent layer (thickness: 300 nm) as detailed in (28). Supplementary Figure 2 shows the spatial intensity maps (Supplementary Fig. 2a, c, e, g) and the average fluorescence intensity along the radial direction (Supplementary Fig. 2b, d, f, h) for uncorrected and corrected type I-IV *eFOV*-microendoscopes. The diameter of the FOV (measured at radial distances at which the measured fluorescence drops below 80%) was significantly higher for corrected compared to uncorrected microendoscopes (356.5 ± 8.4 μm *vs* 218.8 ± 83.4 μm for corrected and uncorrected type I probes, respectively, p = 9E-4, N = 7; 360.3 ± 11.3 μm *vs* 156.9 ± 4.8 μm for corrected and uncorrected type II probes, p = 2E-14, N = 8–9; 163.8 ± 4.5 μm *vs* 131.1 ± 6.0 μm, for corrected and uncorrected type III probes, p = 4E-8, N = 9; 234.3 ± 2.1 μm *vs* 117.3 ± 3.1 μm, for corrected and uncorrected type IV probes, p = 7E-18, N = 8–9. All p-values were obtained with Student’s *t*-test).

We next characterized the effect of aberration correction on the axial resolution across the FOV. From the acquired z-stacks of a subresolution thin fluorescent layer described above, we measured the FWHMz across the FOV. As expected by ray trace simulations (Fig. 1), *eFOV*-microendoscopes displayed a more pronounced curvature of the focal plane (Fig. 3f, Supplementary Fig. 3) compared to uncorrected probes. Moreover, the x,z projection of the z-stacks showed higher axial resolution in *eFOV*-microendoscopes in an area that was ~ 1.2 - 9.3 folds wider (depending on microendoscope type) compared to uncorrected probes (Fig. 3f-g, Supplementary Fig. 3, table 1), further demonstrating extended FOV in corrected microendoscopes. Noteworthy, we found good agreement between the experimental measurements (Fig. 3, table 1) and the prediction of the optical simulations (Fig. 2). Since aberrations generally increase with the length of the GRIN rod used in the microendoscope (38), aberration correction resulted in larger improvements in optical performances in *eFOV*-microendoscopes with longer rather than shorter GRIN rods (Supplementary Fig. 3, table 1). The ability of *eFOV*-microendoscopes to image effectively larger FOV compared to uncorrected probes was further confirmed in biological tissue by imaging neurons expressing the green fluorescence protein (GFP) in fixed brain slices (Fig. 3h, Supplementary Fig. 4).

**Figure 4.**
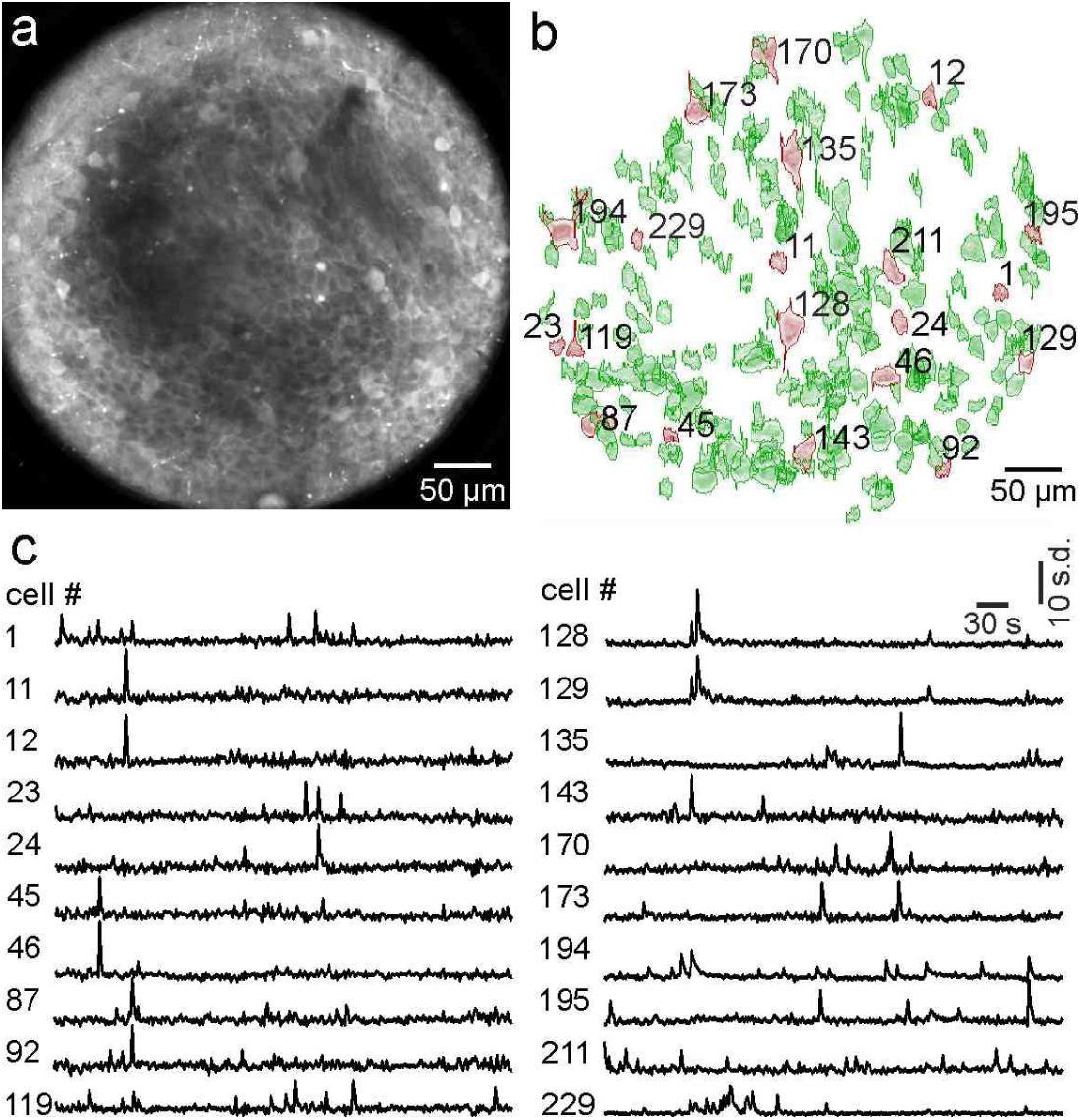
Large FOV functional imaging of hippocampal networks with minimal invasiveness. **a-c)** Two-photon image showing GCaMP6s expressing hippocampal neurons recorded using type I *eFOV*-microendoscopes. Identified active ROIs are shown in b). The fluorescence signal over time for 20 representative ROIs (red in b) are displayed in c). Traces of fluorescence signals for all active ROIs (green in b) are displayed in Supplementary Figure 7.

To validate *eFOV*-microendoscopes performances for functional measurements *in vivo*, we expressed the genetically-encoded calcium indicator GCaMP6s in the mouse hippocampal region (Supplementary Fig. 5a-a_2_) and implanted *eFOV*-microendoscopes above the injected area (Supplementary Fig. 6a, c). It is worth noting that an advantage of the customized mount that we developed to hold *eFOV*-microendoscopes (Fig. 3a-b) is that it allows their full-length insertion within the tissue. As compared to the most common microendoscopes configurations described in the literature (14, 16, 17, 19, 39), our solution thus enables the use of shorter probes, which are less sensitive to aberrations.

We applied *eFOV*-microendoscopes to perform *in vivo* population imaging in injected mice in the raster scanning configuration at 0.5–0.7 Hz (Fig. 4, Supplementary Figs. 7–8). Spontaneous activities in the CA1 hippocampal region were imaged with type I *eFOV*-microendoscopes (Fig. 4, Supplementary Fig. 7) and type III *eFOV*-microendoscopes (Supplementary Fig. 8). Using a cell-sorting algorithm based on PCA/ICA analysis (34), tens to hundreds of active ROIs were identified and could be imaged on a single FOV using *eFOV*-microendoscopes, confirming efficient population imaging (Fig. 4, Supplementary Figs. 7–8). Moreover, neuronal processes and dendritic spines could be reliably monitored in lateral parts of the FOV, demonstrating high-resolution imaging across the whole extended FOV *in vivo* and proving the effectiveness of eFOV-microendoscopes for imaging neuronal activity at both cellular and synaptic level. Longer eFOV-microendoscopes for imaging of deeper brain areas, such as the dorsal striatum and the ventral posteromedial thalamic nucleus (Supplementary Figs. 5–6), were also successfully implanted.

Finally, because the built-in aberration correction method adopted in *eFOV*-microendoscopes does not interfere with the temporal resolution of the optical system, we coupled *eFOV*-microendoscopes with the scanless imaging technique (31, 40) to improve temporal resolution in endoscopic functional imaging. A SLM (Supplementary Fig. 9a) (31–33, 40) was used to generate an array of points in the focal plane, each stimulating a neuron expressing GCaMP6s with a near diffraction limited spot. Simultaneously excited fluorescence at multiple locations was collected through a camera, decoupling the maximal acquisition frequency from the number of imaged points. Using this experimental configuration, we performed simultaneous imaging from multiple hippocampal CA1 pyramidal cells *in vivo* at two orders of magnitude higher speed (125 Hz, Supplementary Fig. 9) compared to scanning imaging (0.5–0.7 Hz, Fig. 4 and Supplementary Figs. 7–8).

Major efforts in the development of technology for imaging neuronal activity *in vivo* are directed to access deep brain regions with minimal tissue damage while maximizing the area over which imaging can be performed with single cell resolution and high sampling speed. GRIN lenses, alone or in combination with fiber-bundles, have been used to perform one- and two-photon imaging in deep brain areas, such as the hippocampus (39, 41), the striatum (16) and the hypothalamus (16, 42, 43). GRIN microendoscopes have also been used to perform simultaneous functional imaging of two different brain regions (44), allowing concurrent monitoring of neuronal dynamics in areas otherwise not accessible with single FOV systems.

Even though the combination of GRIN lenses with two-photon imaging benefits of improved optical sectioning, most GRIN endoscopes have been operating with lower resolution even in two-photon excitation modality due to optical aberrations lying on and off the optical axis. These aberrations derive from the intrinsic nonaplanatic properties of GRIN rods (45) and limit the usable FOV (14, 17, 19, 21). Aberrations increase with the length and NA of the GRIN lens, making high-resolution imaging of large FOV in deep areas a challenging goal. Aberrations can be partially compensated by adding additional optical elements, such as a cover glass (38), a single high refractive index planoconvex lens (14), and multiple planoconvex lenses combined with diffractive optical elements (21). By adding a single planoconvex ball lens to the distal end of a customized GRIN rod (rod diameter: 1 mm), Barretto et al. increased the NA up to 0.82 correcting on-axis aberrations and they imaged neuronal dendritic spines in GFP-expressing hippocampal pyramidal neurons in live mice (14). To correct off-axis aberrations while maintaining high NA, at least a second planoconvex lens was needed (21). Fabrication of high performances multi-element optical systems for on-axis and off-axis aberration correction, however, imposes to respect strict tolerances in the assembly and needs the use of external metal cannulas to hold the various optical elements aligned and provide mechanical stability to the optical system. This, so far, limited the applicability of aberration correction with built-in optical elements to GRIN lenses of diameter ≥ 1 mm and overall endoscopic probe diameter (GRIN + cannula) of ≥ 1.4 mm (14, 21). Since the insertion of the probe irreversibly damages the tissue above the target area, reducing the size of the probe and consequently its invasiveness is of outmost importance when imaging deep brain regions. However, due to their small radial dimensions, improving optical performances in ultrathin (diameter ≤ 0.5 mm) microendoscopes with built-in optical elements is a major challenge. In this study, we devised a new approach to solve this problem and used TPP (22, 46) to microfabricate polymeric aspheric lenses that effectively corrected aberrations in ultrathin GRIN-based endoscopes. Corrective lenses were first fabricated on glass coverslips which were aligned and assembled with the GRIN rod to form an aberration-corrected microendoscope. Importantly, this optical design resulted in improved axial resolution and extended FOV without increasing the lateral size of the probe and thus minimizing tissue damage in biological applications.

Aberration correction in GRIN microedoscopes can be achieved using adaptive optics (AO) (17, 19, 20, 45). For example, using pupil-segmentation methods for AO diffraction-limited performance across an enlarged FOV was obtained in GRIN-based endoscopes with diameter of 1.4 mm (17, 19) and, in principle, this approach could be extended to probes with smaller diameter. AO through pupil segmentation requires significant modification of the optical setup and the use of an active wavefront modulation system (e.g. deformable mirror device or liquid crystal spatial light modulator) which needs the development of ad-hoc software control. Moreover, AO through pupil segmentation may limit the temporal resolution of the system, since multiple AO corrective patterns must be applied to obtain an aberration-corrected extended FOV (17). Compared to AO approaches, the technique developed in this study does not necessitate modification of the optical path nor the development of heavy computational approaches. Moreover, it is easily coupled to standard two-photon set ups, and does not introduce limitations in the temporal resolution of the imaging system, allowing fast endoscopic microscopy when coupled with the scanless imaging modality (Supplementary Fig. 9).

Despite these advantages, the approach presented in this study is limited to the correction of aberration introduced by the GRIN lens and was not developed to correct aberrations introduced by other elements along the optical path or by the sample. Moreover, in contrast to AO our approach does not correct for aberrations that may vary over time, such as those due to intrinsic properties of the biological sample that may dynamically change over the course of the experiment. AO approaches could, for example, be applied on *eFOV*-microendoscopes to address this limitation. Future development of the method described in this study may also include the realization of corrective elements composed of compound microfabricated lenses (22, 46) that could extend the degrees of freedom in the correction design process and lead to improved performances of miniaturized optical probes.

## Conclusions

We developed a new methodology to correct for aberrations and extend the FOV in ultrathin microendoscopes using microfabricated aspheric lenses. Corrective lenses are specifically designed for each type of GRIN rod used in the endoscope and are fabricated by TPP. This method is flexible and can be applied to the GRIN rods of different diameters and lengths that are required to access the numerous deep regions of the mammalian brain. Corrected endoscopes showed improved axial resolution and up to 9 folds extended FOV, allowing efficient *in vivo* population imaging with minimally invasiveness. Moreover, *eFOV*-microendoscopes can be efficiently coupled to fast imaging approaches to increase the temporal resolution of aberration-corrected endoscopic imaging by almost two orders of magnitude.

Although *eFOV*-microendoscopes have been primarily applied for functional imaging in this study, we expect that their use can be extended to other applications. For example, *eFOV-*microendoscopes could be combined with optical systems for two-photon patterned optogenetic manipulations (47–49) and for simultaneous functional imaging and optogenetic perturbation (50–52). Moreover, besides its applications in the neuroscience field, *eFOV*-microendoscopes can be used in a large variety of optical applications requiring minimally invasive probes, ranging from cellular imaging (35, 53) to tissue diagnostic (54, 55). Importantly, applications of ultrathin *eFOV-*microendoscopes to other fields of research will be greatly facilitated by the built-in aberration correction method that we developed. This provides a unique degree of flexibility that allows using ready-to-use devices in a large variety existing optical systems with no major modification of their optical path.

## Acknowledgements

We thank M. Dal Maschio for discussion at an initial stage of the project, B. Sabatini and A. Begue for critical reading, V. Jayaraman, J. Akerboom, R. A. Kerr, D. S. Kim, L. L. Looger, K. Svoboda for GCaMP. This work was supported by an IIT interdisciplinary grant to TF and CL and in part by ERC (NEURO-PATTERNS), FP7 (DESIRE), MIUR FIRB (RBAP11X42L), and Flag-Era JTC Human Brain Project (SLOW-DYN) to TF.

## Author contributions

AA, SB, CM and FS performed experiments and analysis. AA performed simulations and microendoscope fabrication. AB performed microendoscope fabrication with the Nanoscribe system. CM developed hardware and software for patterned illumination. VPR developed polymeric materials and molding replication process. TF and CL conceived and coordinated the project. All authors contributed to writing and approved the final version of the manuscript.

## SUPPLEMENTARY MATERIAL

**Supplementary figure 1.**
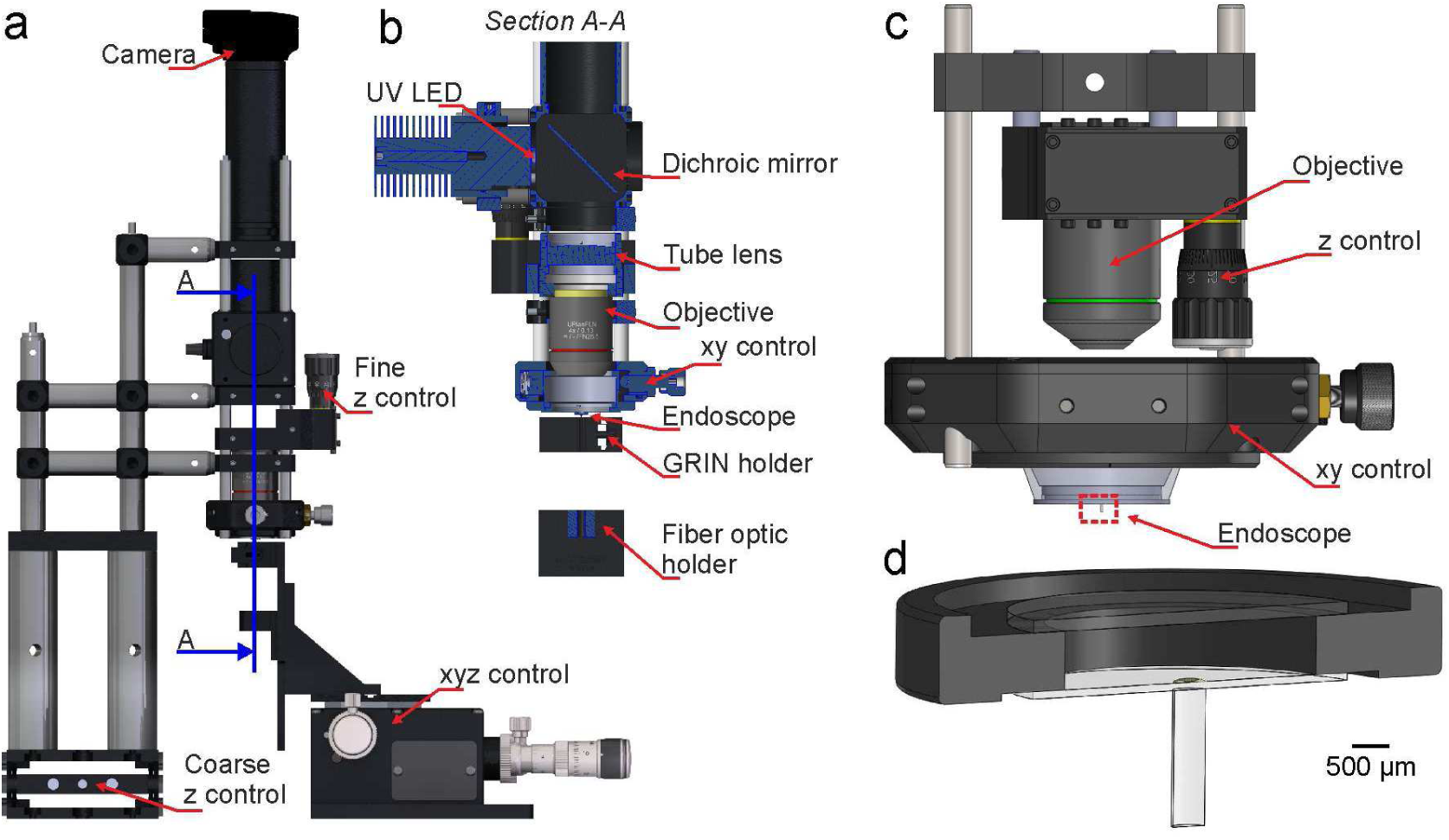
Set ups for the assembly and characterization of *eFOV-*microendoscopes. **a)** Optomechanical stage used for microendoscope assembly. Arrows indicate key components: camera (DCC1645C), fine z control (SM1Z), coarse z control (L200/M), and xyz control (MAX313D/M). All items were purchased from Thorlabs. The blue line indicates the part of the set up whose section is shown at an expanded scale in b). **b)** Section of a portion (A-A) of the set up shown in c). Key components were: high power UV LED (M375L3, Thorlabs), long pass dichroic mirror (FF409-Di02, Semrock), tube lens (AC254-150-A, Thorlabs), objective (UPlanFLN 4x 0.13NA, Olympus), xy control (CXY1, Thorlabs), custom GRIN rod holder, fiber optic holder (HCS004, Thorlabs). **c)** Schematic of the optomechanical assembly used for microendoscopic imaging. The coupling objectives were RMS20X-PF-20X and LUCPLFLN 60X (Olympus). The z control (SM1Z) and xy control (CXY2) were purchased from Thorlabs. **d)** The self-supported *eFOV*-microendoscope (highlighted with the red dotted line in c) is shown at an expanded scale.

**Supplementary figure 2.**
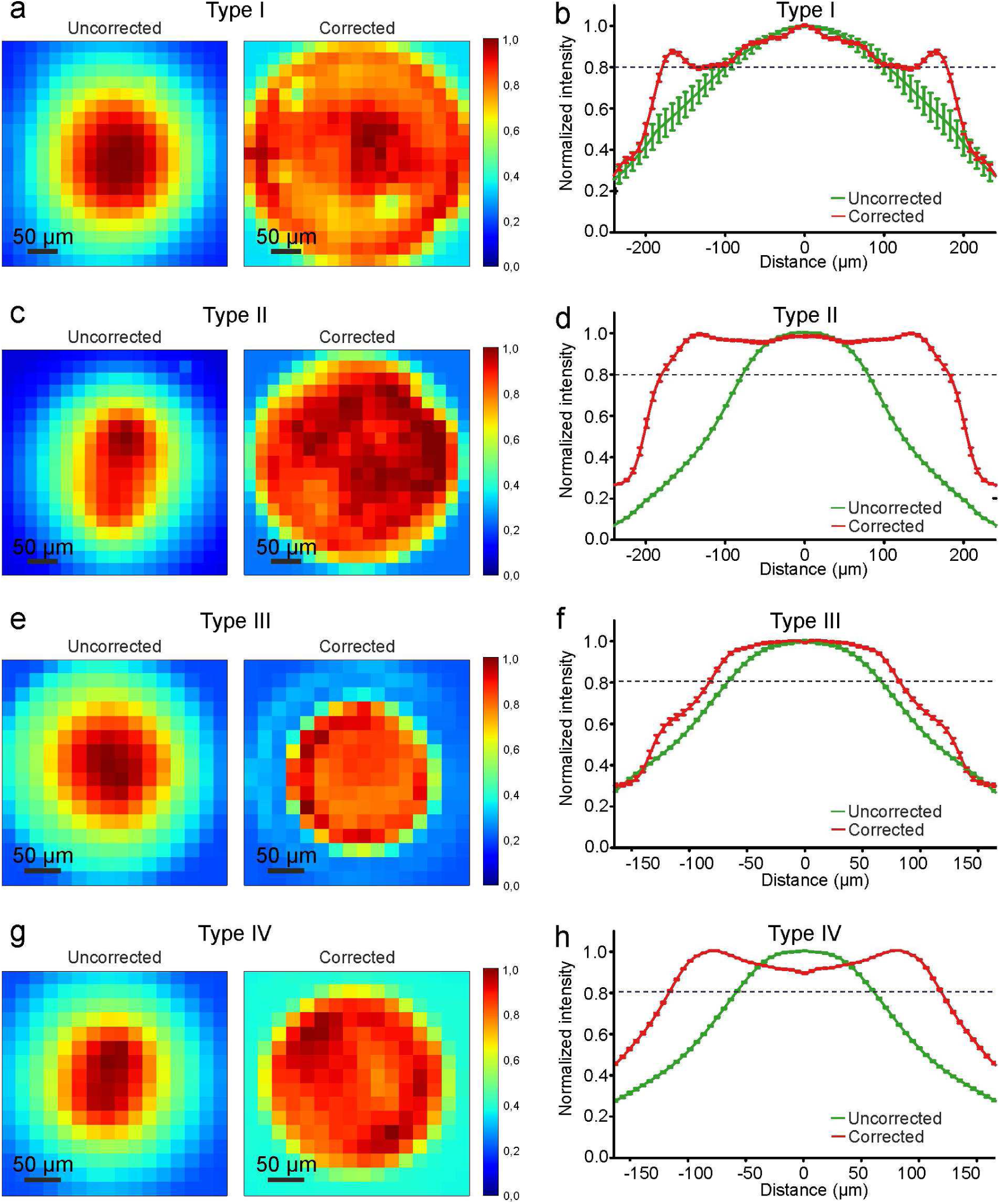
Aberration correction increases the FOV. **a)** Pseudocolor images showing the spatial distribution of fluorescence of uncorrected (left image) and corrected (right image) type I microendoscopes. Images are obtained monitoring a thin homogeneous fluorescence layer at λ_exc_ = 920 nm and were resampled (pixel dimension: 20 μm) to suppress sharp fluorescence variations. **b)** Fluorescence intensity profile along a line crossing the optical axis for uncorrected (green line) and corrected (red line) type I microendoscopes. Traces are shown as mean ± sem across different experiments (N = 7). **c-d)** Same as in a-b) for type II microendoscopes. In d), N = 8–9. **e-f)** Same as in a-b) for type III microendoscopes. In f), N = 9. **g-h)** Same as in a-b) for type IV microendoscopes. In h), N = 8–9.

**Supplementary figure 3.**
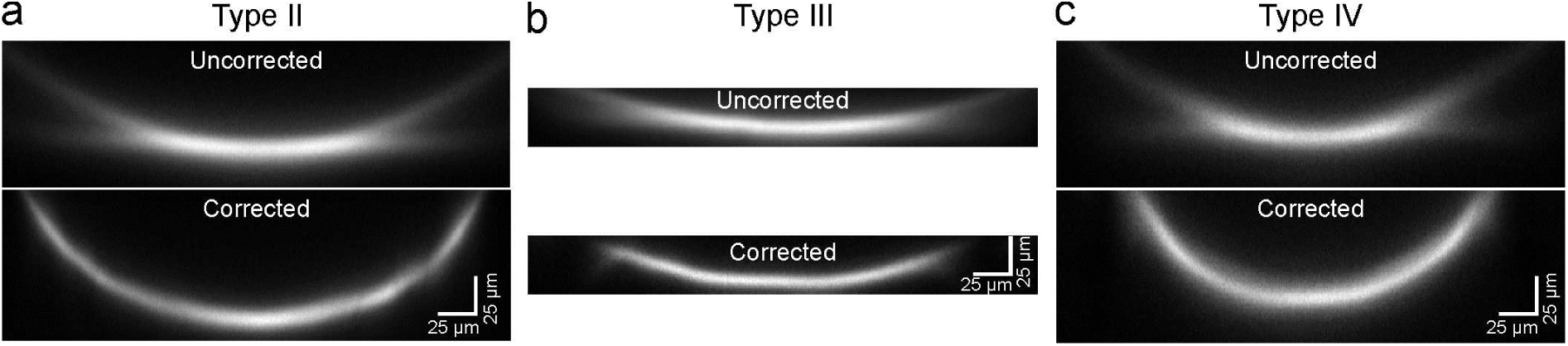
Aberration correction with polymeric lenses increases the FOV in GRIN-based microendoscopes. **a-c)** x,z projections of a z-stack of two-photon laser scanning images of a subresolution fluorescent layer (thickness: 300 nm) without (uncorrected, top) and with (corrected, bottom) corrective lens for type II (a), type III (b) and type IV (c) *eFOV*-microendoscopes. Excitation wavelength: λ = 920 nm.

**Supplementary figure 4.**
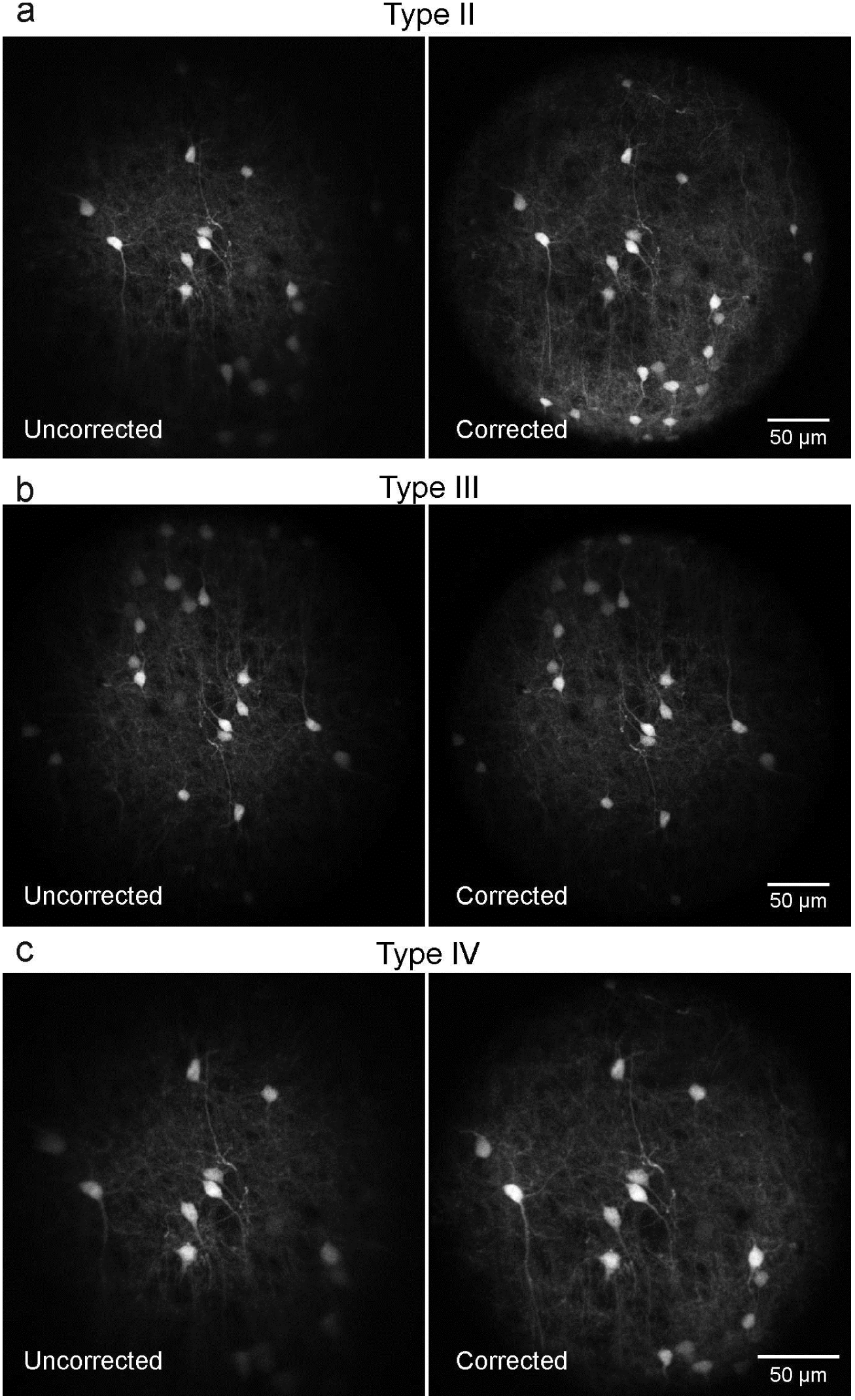
Extended FOV in corrected microendoscopes. **a-c)** Representative images of fixed cortical tissue expressing GFP in neuronal cells were acquired with type II (a), type III (b) and type IV (c) *eFOV*-microendoscopes without (uncorrected, left panels) and with (corrected, right panels) corrective lens.

**Supplementary figure 5.**
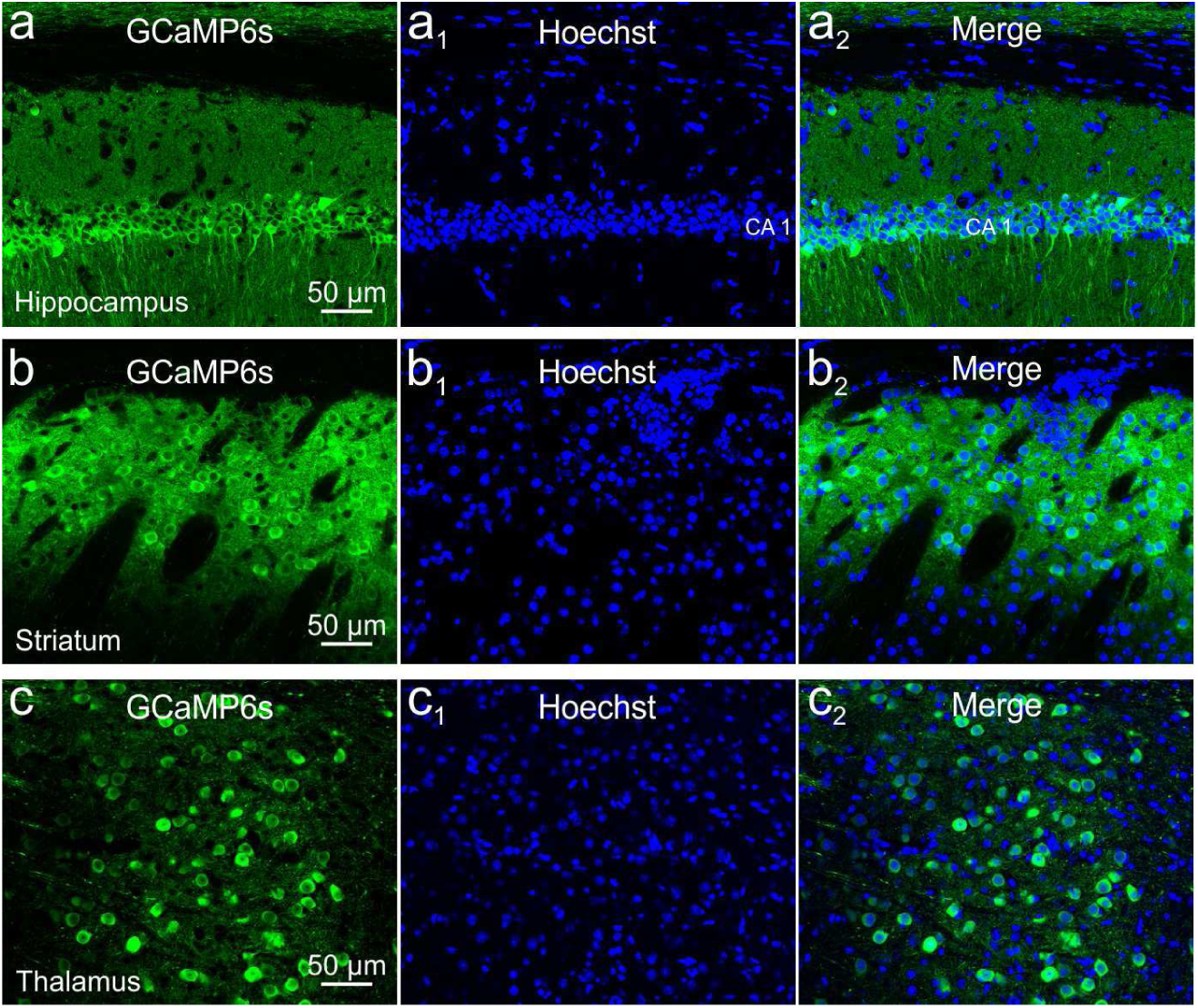
GCaMP6s expression in deep brain areas. **a-a_2_)** Confocal images of hippocampal CA1 neurons expressing GCaMP6s (a). Nuclei were counterstained with Hoechst (a_1_). Images are merged in a_2_. Scale bar in a applies to a_1_-a_2_. **b-b_2_**) Same as in a-a_2_ for neurons in the dorsal striatum. **c-c_2_**) Same as in a-a_2_ for neurons in the ventral posteromedial thalamic nucleus.

**Supplementary figure 6.**
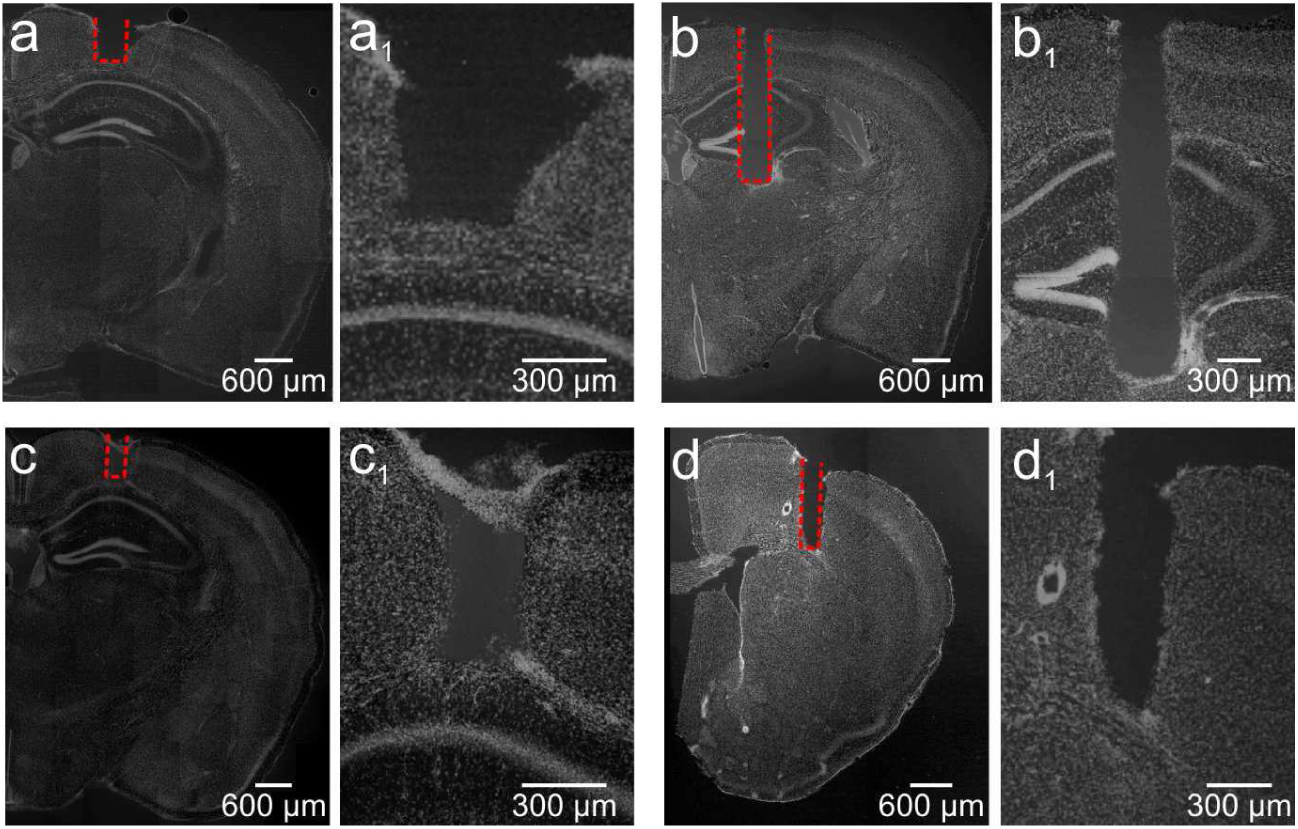
Implantation of different types of *eFOV*-microendoscopes in various brain areas. **a-d_1_)** Confocal images showing coronal slices from mice implanted with type I (a-a_1_), type II (b-b_1_), type III (c-c_1_) and type IV (d-d_1_) *eFOV*-microendoscopes. Type I and III *eFOV-*microendoscopes were used to reach the hippocampus, type II to reach the thalamus and type IV to reach the striatum. Slices were counterstained with Hoechst. The probe track is highlighted with the red dotted line in a, b, c, d and shown at a higher magnification in a_1_, b_1_, c_1_, d_1_.

**Supplementary figure 7.**
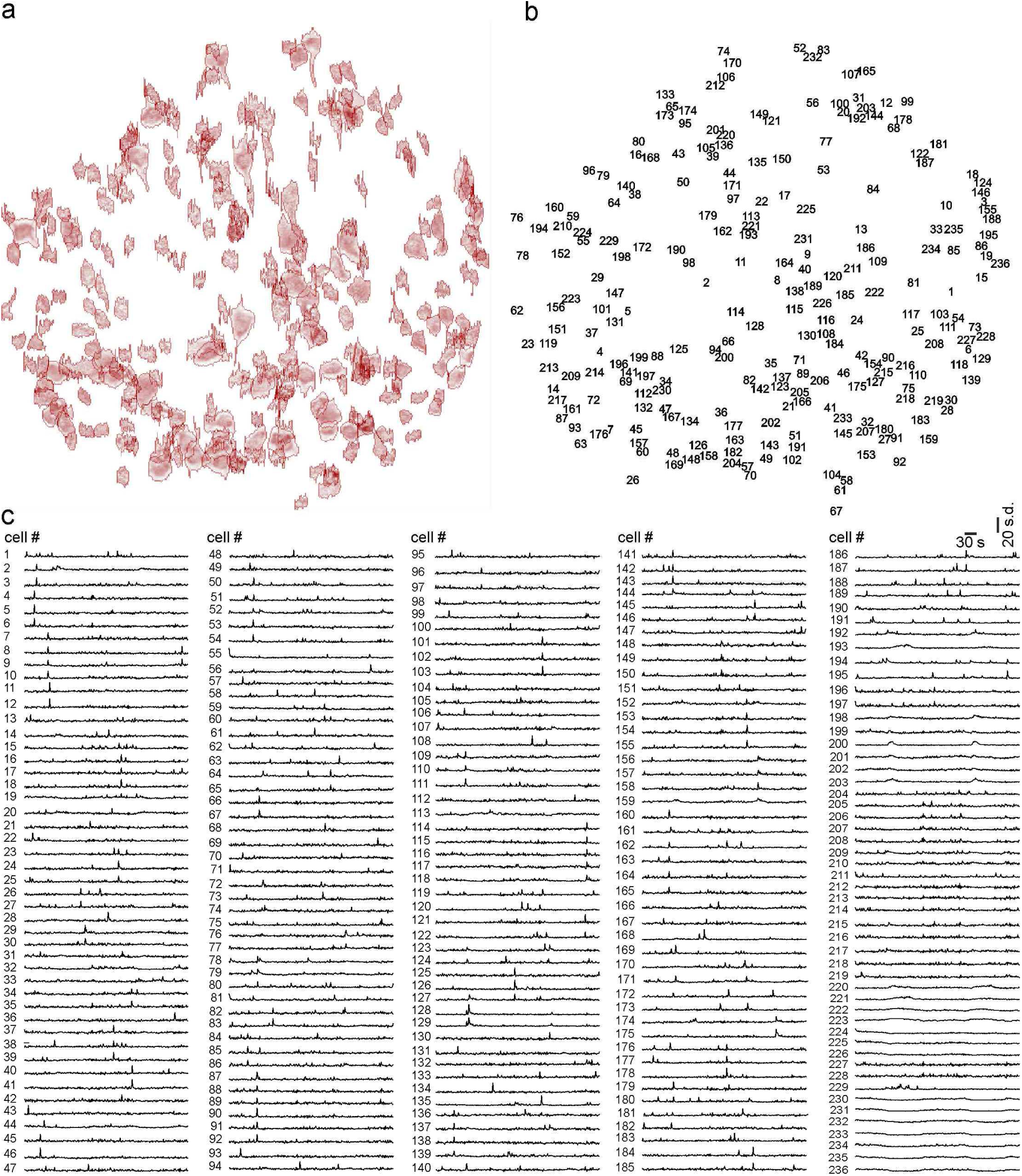
Population imaging with ultrathin type I *eFOV*-microendoscopes. **a-b)** Active ROIs are identified (a) and numbered (b) for the hippocampal field displayed in figure 4a. **c)** Fluorescence signals over time for the ROIs displayed in a-b).

**Supplementary figure 8.**
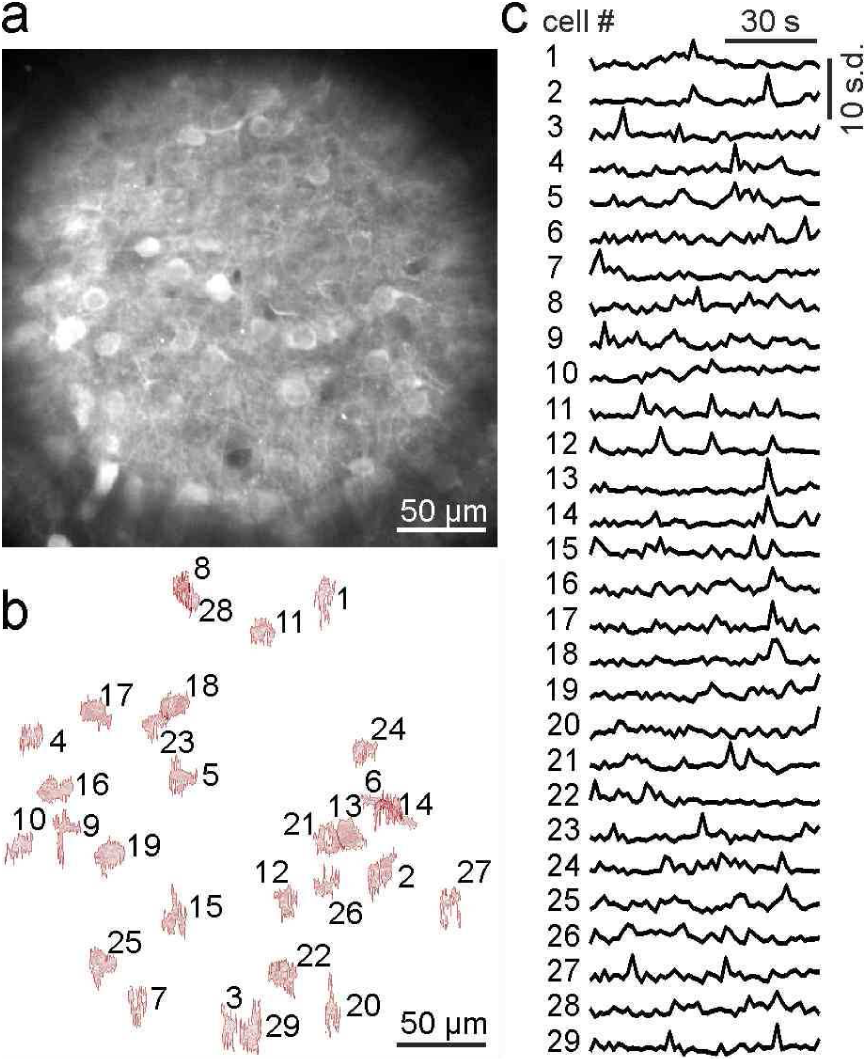
Population imaging with ultrathin type III *eFOV*-microendoscopes. **a-b)** GCaMP6s expressing hippocampal neurons. Active ROIs are identified and numbered in b). **c)** Fluorescence signals over time for the ROIs displayed in **b)**.

**Supplementary figure 9.**
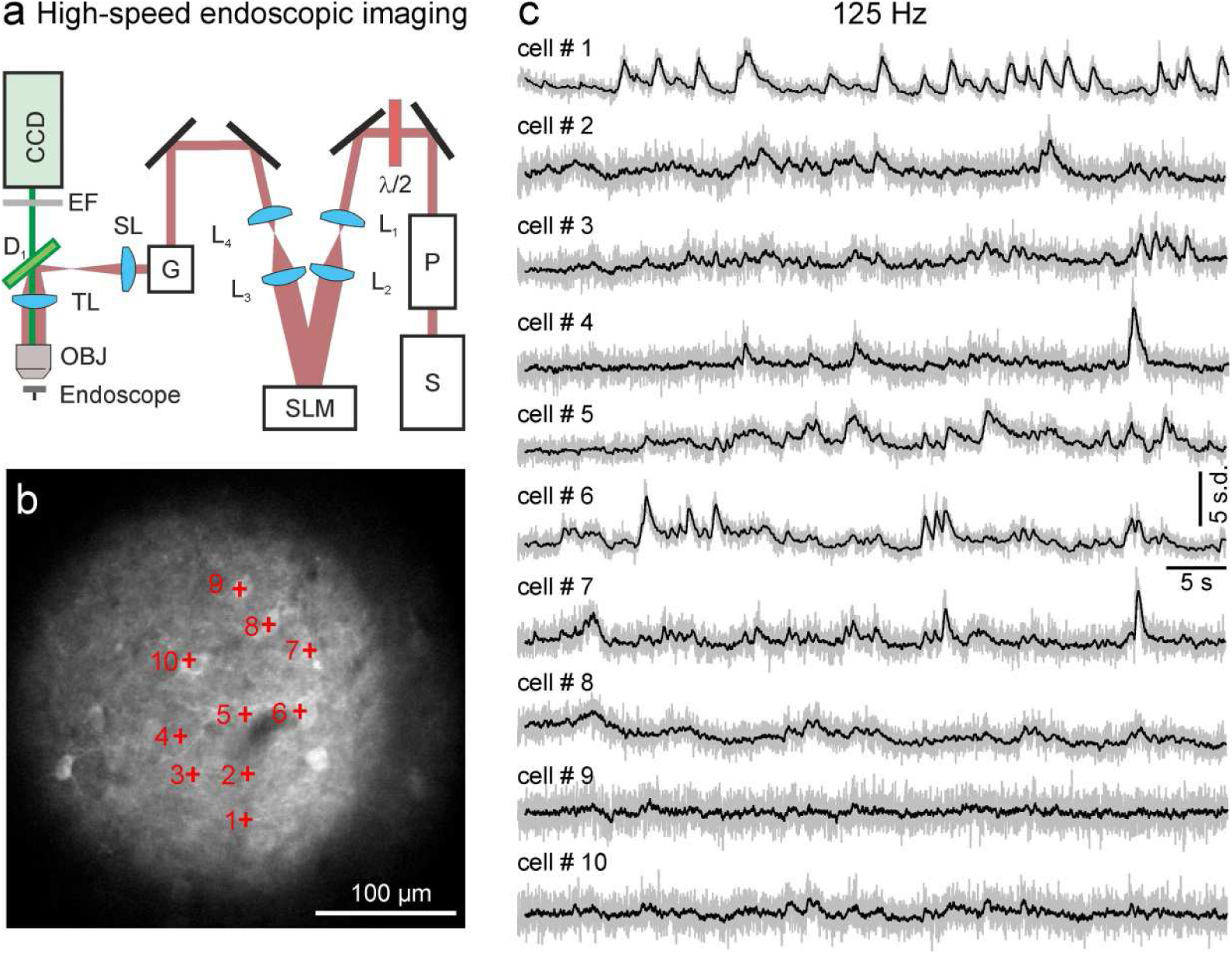
High-speed functional imaging with *eFOV*-microendoscopes. **a)** Schematic of the experimental set-up for fast scanless microendoscopic imaging. A SLM is used to spatially multiplex the laser beam and fluorescence signals are collected via a camera. A custom code was written in LabView (National Instruments Corp, Austin, TX) to compute phase modulation maps and to interface the SLM with the PrairieView acquisition software (Bruker Corporation, Milan, IT). Excitation patterns with arrays of points were generated using the Weighted Gerchberg-Saxton Iterative Fourier Transform Algorithm. A calibration routine with submicrometric precision was developed to map the FOV of the laser scanning system with the projection plane of the SLM at the sample. The light of the zero-order diffraction component was typically projected to a region where non-fluorescent structures were identified (e.g.: the lumen of a blood vessel). **b)** Two-photon laser scanning image showing GCaMP6 expressing neurons in the CA1 hippocampal region. A type III *eFOV*-microendoscope was used. Labelled neurons (red crosses) were imaged in the scanless configuration. **c)** Fluorescence signals over time for the neurons displayed in b). Raw traces are shown in grey; black lines represent a posteriori filtered traces (see Materials and Methods).

**Supplementary table 1.** Coefficients used in equation 1 (see Materials and Methods) for the aspherical corrective lenses used in type I-IV *eFOV*-microendoscopes.

|  | **c** | **k** | **$\alpha_1$** | **$\alpha_2$** | **$\alpha_3$** | **$\alpha_4$** | **$\alpha_5$** | **$\alpha_6$** | **$\alpha_7$** | **$\alpha_8$** |
| --- | --- | --- | --- | --- | --- | --- | --- | --- | --- | --- |
| **Type I** | -2.58E-01 | -1.74E+00 | 8.58E-01 | -5.30E+01 | 5.95E+03 | -2.77E+05 | 7.26E+06 | -8.91E+07 | 2.47E+08 | 2.19E+09 |
| **Type II** | 1.02E+03 | -2.37E+03 | -1.46E+00 | 2.87E+01 | -4.60E+01 | -7.69E+03 | 1.96E+05 | 7.82E+06 | -1.69E+08 | -3.88E+08 |
| **Type III** | -4.99E-01 | 8.20E+00 | 1.54E+00 | -1.35E+02 | 3.67E+04 | -3.92E+06 | 2.17E+08 | -5.56E+09 | 3.34E+10 | 6.29E+11 |
| **Type IV** | -2.41E-01 | -1.17E+00 | 9.22E-01 | -1.34E+02 | 4.36E+04 | -4.28E+06 | 2.23E+08 | -5.55E+09 | 3.48E+10 | 4.94E+11 |

